# Parallelised detection of bacteria viability using an electrode array and the Exeter Multiscope

**DOI:** 10.64898/2026.03.10.710830

**Authors:** Ka Kiu Lee, David Horsell, James Stratford, Magdalena Karlikowska, Salman Khattak, Tailise de-Souza-Guerreiro-Rodrigues, Junqing Jiang, Mike Shaw, Stefano Pagliara, Alexander D. Corbett

## Abstract

Antimicrobial resistance remains a global existential threat. Given that antimicrobial therapy commonly starts before pathogen identification, rapid and scalable methods capable of determining effective antimicrobial compounds are needed. In this paper, we demonstrate a 2 × 2 array of parallelised microscopes that uses low numerical aperture (NA=0.25) detection optics and LED excitation to determine bacterial viability based on their fluorescence response to an electrical stimulus. Following a 2-hour incubation, the fluorescent viability readout requires less than one minute. We use K-means clustering to classify pixels in a time lapse sequence of widefield fluorescence images and extract changes seen within bacterial clusters. We demonstrate sufficient sensitivity to measure fluorescence changes after electrical stimulation in a bacterial monolayer. To capture these subtle fluorescence changes at high signal-to-background ratios, we place a limit on the minimum optical density of the bacterial sample. This novel approach is scalable to 96-well formats using a suitable consumable electrode array.

## Introduction

Antimicrobial resistance (AMR) remains a major health threat to the global population. The annual cost of AMR in Europe alone is currently estimated at €1.5 bn with AMR predicted to cause 10 million deaths annually by 2050^1^. Developing a rapid and scalable AMR test would reduce this burden by ensuring patients receive the appropriate antibiotic at the right concentration sooner.

The current clinical gold standard for determining antibiotic resistance in a patient sample is to culture the bacterial isolate with a panel of antibiotics across multiple concentrations for 24 to 48 hours. The efficacy of an antibiotic and its minimum inhibitory concentration (MIC) are established by monitoring bacterial growth (turbidity) in in each well. The limitation of this approach is that, in acute cases, results are required within much shorter time frames (a few hours) for effective treatment. The long incubation period also limits throughput and contributes to a backlog of testing.

In recent years, there has been a rapid expansion of antimicrobial susceptibility testing (AST) methods available (see Reszetnik, et al. [^2^] for a review). Recent advances include direct-from-specimen testing, single-cell imaging, miniaturized growth chambers, microfluidics, and novel detection modalities including plasmonic and nucleic acid-based methods that shorten turnaround times by accelerating growth detection or viability assessment under antimicrobial exposure. Several commercial platforms now provide phenotypic AST directly from positive blood cultures or urine specimens significantly faster than traditional methods (such as the electric field detection method employed by iFast Diagnostics^3^). Emerging non-commercial approaches show promise for further reductions in time-to-result, including methods that combine rapid imaging or metabolic sensing with deep learning for enhanced signal interpretation. Challenges remain in extending these technologies to diverse specimen types, standardising inoculum effects, and achieving broad clinical validation.

One recent technique developed by the Munehiro lab^4^ measures bacterial viability via electrical stimulation of the bacteria under investigation, whilst monitoring changes in fluorescence intensity, with positive and negative changes indicating viable and non-viable bacteria, respectively. After incubation in a medium containing fluorescent dye thioflavin T (ThT), electrical stimulation hyperpolarises the membrane of healthy cells by opening voltage-sensitive ion channels. Viable bacteria rapidly accumulate surrounding (positively charged) ThT through these ion channels, leading to increased fluorescence brightness. In contrast, non-viable bacteria with disrupted membranes and a weakened potassium ion gradient will undergo depolarisation and expel any intracellular ThT through electrical stimulation. This translates into either no change or a reduction in fluorescence brightness.

This technique has since been refined and commercialised^5^. Whilst significant developments have been made to improve sensitivity and repeatability, there is clear potential (as with many of the techniques summarised in [^2^]) to greatly increase sample throughput. This could be achieved by creating arrays of stimulation electrodes, each with their own bacterial sample. However, in studies performing fluorescence imaging of multiple fields of view, 75% of the total acquisition time was spent moving the sample^6^, with similar estimates for fast axial scanning systems^7^. Throughput can be dramatically increased by eliminating the need to move samples by combining electrical stimulation with parallelised microscopy.

Parallelised microscopy, in which an array of samples can be imaged individually without the need to mechanically translate the sample, has gained popularity in recent years. Unlike macroscopic imaging of the entire sample array, where each sample occupies a small fraction of the whole image, in parallel microscopy, each sample is imaged by the entire detector area, greatly increasing the achievable resolution from ∼100 µm to less than 10 µm. Versions of parallelised microscopes include schemes that have a separate excitation and detection path for each sample, using arrays of light sources, lenses, and cameras to capture image data from each sample simultaneously^8,9^. This approach has the advantage of true simultaneity for up to 24 samples, but the machine vision cameras used lack the sensitivity required to detect the fluorescent emission from small bacterial microcolonies, which may be less than 10 µm across. In addition, the large data rates associated with collecting data from 24 cameras at once (45 Gbps compared to 10 Gbps for USB 3.1 Gen 2.0) require custom data transfer methods, limiting scalability and impact. An alternative method is to use a galvanometer mirror to sweep a macroscopic image of all samples across a single detector to increase sampling at the desired locations, but this low N.A. (i.e. 0.04) imaging method lacks spatial resolution^10^.

“Random Access Parallel” (RAP) microscopy^11^ uses an array of LED light sources to illuminate each sample individually, but uses a common detection path and therefore requires only one camera. This sequential acquisition approach has recently been applied to the measurement of cardiomyocyte monolayer contraction (implemented as the “Exeter Multiscope”)^12^. The utility of using one detection camera becomes particularly apparent in fluorescence imaging, as the prospect of having an array of low-noise, high quantum efficiency fluorescence cameras quickly becomes impractical in terms of mechanical integration and cost. We have developed a proof-of-concept parallelised microscope with the ability to distinguish between viable and non-viable bacteria using a custom-built UV-fluorescence Multiscope that can read out a 2 × 2 array of bacterial samples, testing up to four different conditions, in rapid succession, without any moving parts in less than 5 minutes. K-means clustering is used to extract fluorescence intensity changes associated with bacterial clusters from time lapse data. We demonstrate the fluorescence detection sensitivity required to scale the readout to larger numbers of bacterial samples, including standard 96-well plate formats.

## Results

### Baseline results from an inverted microscope

To provide a baseline measurement of bacterial fluorescence changes in response to electrical stimulation, we first performed the viable/non-viable assay on an inverted microscope using a multi-electrode array composed of gold electrodes on a coverglass-bottom dish (Figure 1A-B). The electrodes were stimulated individually (100 Hz, 4V peak-peak, 2.5 seconds) using an external control box (‘CytePulse’, Cytecom Ltd.) connected to the electrode array via an IDE ribbon cable. Two of the six electrodes were used to collect reference data (electrode #5 and #6, Figure 1B).

**Figure 1.**
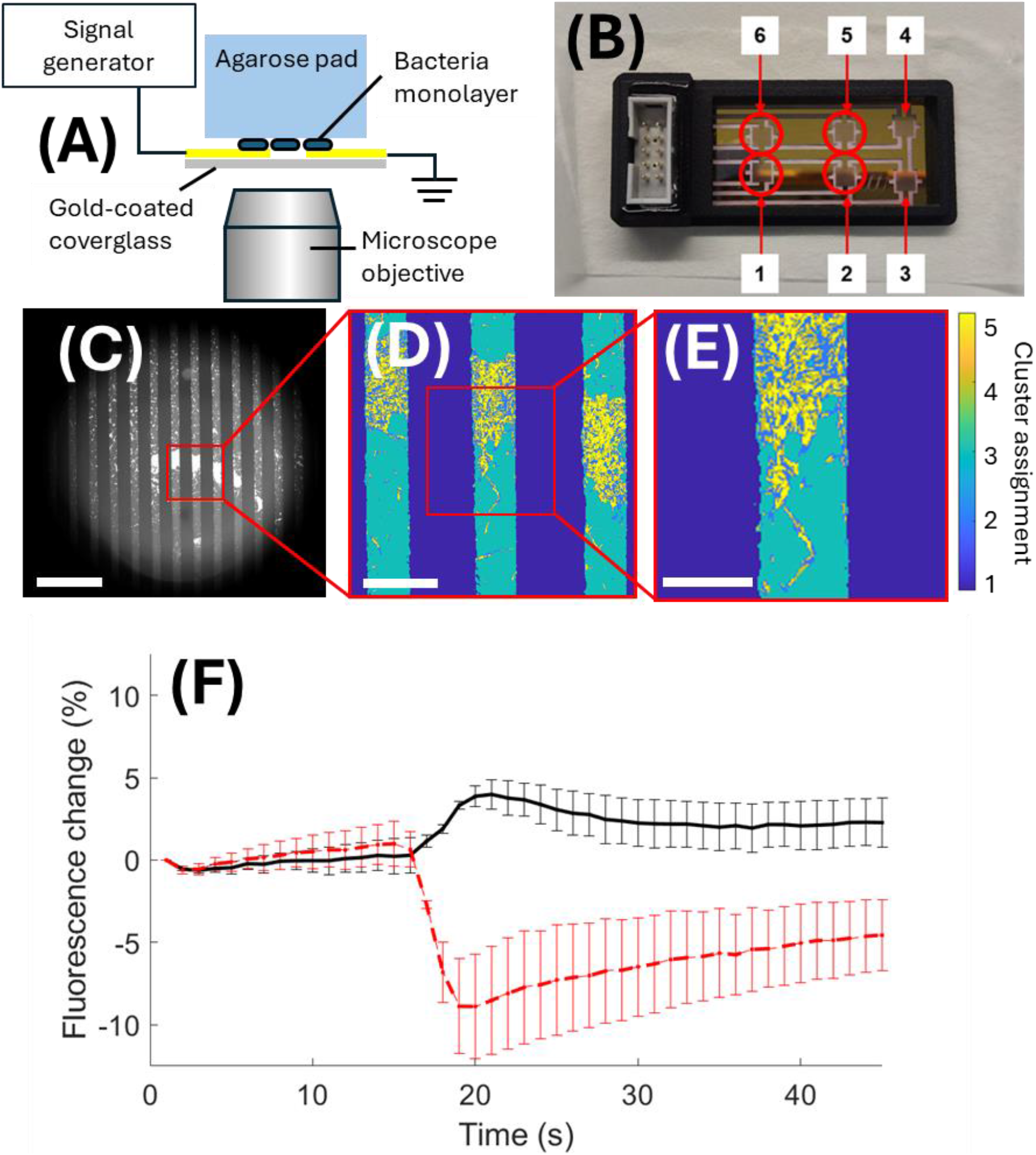
(A) Overview of the method used to electrically stimulate bacterial samples in an inverted microscope. (B) Plan view of an array of six electrode positions. Only positions #5 and #6 were used. (C) Widefield fluorescence image of a bacterial colony, imaged through the electrodes (black vertical bars) on an inverted microscope. One region of interest (ROI) was selected from within this larger field of view for processing. (D) Cluster assignment maps for pixels within each ROI. The top cluster (most strongly correlated to bacterial clusters) was used to calculate the fluorescence change after stimulation. (E) Detail of cluster assignment map indicates the degree of heterogeneity within each colony of B. subtilis. (F) Changes in fluorescence observed after first stimulation (viable bacteria average response, black line), and second stimulation (non-viable bacteria average response, red dashed line). Error bars indicate standard deviation (N=2). Scale bars: (C) 500 µm (D) 100 µm (E) 50 µm.

To measure bacterial viability, *Bacillus subtilis* was cultured to log-phase and an optical density (OD) of 1.5. 1 μL of this culture was pipetted onto one side of 5 × 5 mm agarose pads containing ThT. The inoculated side of each agarose pad was then placed in direct contact with the individual electrodes in the array, ensuring close contact. The pads were then incubated at room temperature for 2 hours to allow bacterial proliferation.

After loading the electrode array onto the microscope stage, a widefield fluorescence image of each sample (electrodes #5 and #6) was taken to confirm the presence of bacteria colonies (Figure 1C). Fluorescence time lapse images were acquired for each pad in sequence using a 10× 0.4 N.A. objective lens and a UV LED source (400 nm, p300 ultra, CoolLED Ltd.). Each time lapse acquired 15 seconds of images (1 image every second, 500ms exposure) before stimulation to provide a fluorescence reference value and then for a further 30 seconds after stimulation to monitor the change (Figure 1F).

The first bacterial assay used proliferating, healthy *B. subtilis* cells. As the process of electrical stimulation is detrimental to the health of the cell membrane, the best measure of the non-viable cell assay was to repeat the experiment on the same site whilst monitoring changes in cell fluorescence after a second stimulation. This repeatable approach avoids complications associated with antibiotic-based killing, such as incomplete loss of viability, while enabling interrogation of the same cell population on the same pad.

To map out the temporal variations in fluorescence intensity over the field of view, a K-means clustering approach was taken. The temporal variation in fluorescence intensity was extracted for each individual pixel in the 256 × 256 pixel region of interest (i.e. over 65,000 fluorescence traces). The K-means clustering algorithm then identified five fluorescence traces (or ‘clusters’) that best characterised the majority of traces present (in a way not dissimilar to principal components analysis). The clusters were ranked according to the magnitude of fluorescence change, with the largest changes being the ‘top’ cluster (=5) and the smallest changes being the ‘bottom’ cluster (=1). The images in Figure 1D and Figure 1E have been colour coded according to the cluster to which each pixel has been assigned. A total of five clusters were chosen to ensure that there were enough to describe the key features anticipated in each field of view: bacteria in the focal plane, bacteria just beyond the focal plane, agarose pad, electrodes, plus an additional cluster for any unanticipated features. Specifying substantially more than five clusters would result in redundant classifications with highly similar cluster characteristics. The clusters were not constrained to be equally populated, as the bacterial clusters occupy a relatively small fraction of the total pixel number.

After clustering, a heat map was produced describing the cluster assignment for each pixel (Figure 1D-E). The cluster that was most strongly associated with the in-focus bacteria (‘top cluster’) was used as the best measure of the change in fluorescence intensity over time (Figure 1F). The absolute fluorescence change was obtained from this cluster by calculating the peak fluorescence difference between stimulation and end points. The peak change in fluorescence for electrode positions #5 and #6 are recorded in Table 1. Table 1 also records the mean pixel value of the traces for the top and bottom cluster from which the signal to background ratio is calculated.

**Table 1:** Summary of fluorescence changes detected using the inverted microscope. Bacteria at each electrode position were stimulated twice before moving to the next electrode.

| Electrode position | Stimulation number | Peak fluorescence change ($\Delta f/f$) | Top trace avg. (a.u.) | Bottom trace avg. (a.u.) | Signal to background |
| --- | --- | --- | --- | --- | --- |
| 5 | 1 | +4.9% | 622.3 | 4.4 | 141.4 |
| 5 | 2 | -12.0% | 585.4 | 4.3 | 136.1 |
| 6 | 1 | +3.2% | 449.8 | 3.8 | 118.4 |
| 6 | 2 | -6.0% | 331.6 | 4.0 | 82.9 |

### Detection of bacterial viability using the Multiscope

The RAP/Multiscope approach to parallelised microscopy is outlined in Figure 2. The samples are arranged on a 9 mm pitch (as in a 96 well plate) and are indicated by the electrode positions in Figure 1B. Each sample is illuminated by collimated light from a UV LED. Baffles reduce cross-talk between each of the illumination paths, ensuring each LED only illuminates one sample. An objective lens below each sample collects emitted light to form an image at infinity. The final element in this system has a diameter large enough to cover all of the samples in the array and acts to both steer and focus the light from each sample to a common exit pupil, which is in the same plane as the camera. This final element could be a transmissive lens, but for the purposes of maintaining a compact setup, a reflective parabolic mirror was used. To maintain the camera at the focus of the parabolic mirror, whilst not obscuring the light from the sample, the axis of the mirror was offset from the centre of the array. A photograph of the prototype system used in this project is shown in Figure 2(B).

**Figure 2.**
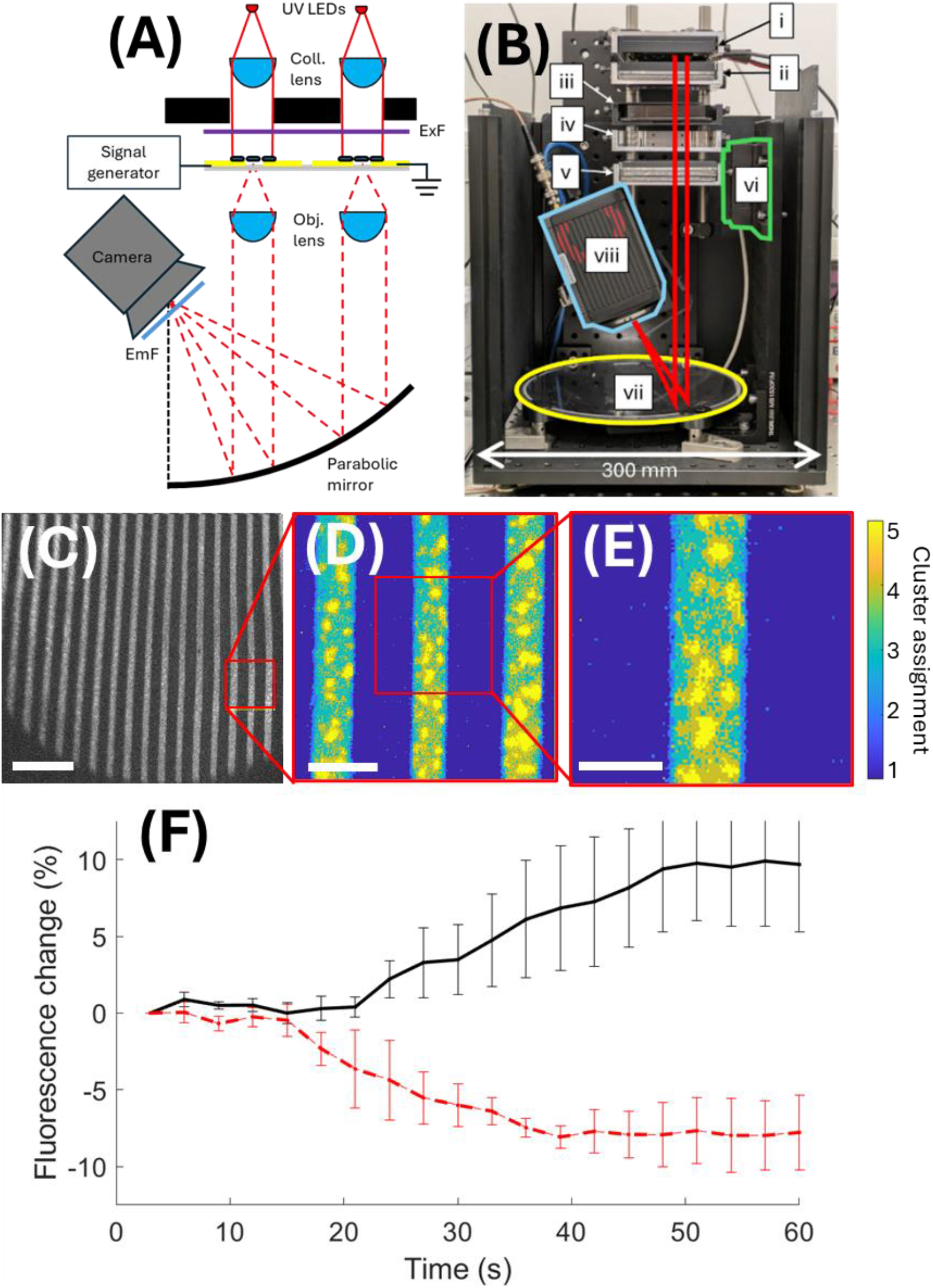
(A) Overview of the method used by the Multiscope to parallelise imaging of multiple samples. Each sample is illuminated one at a time and the images are relayed to the same sCMOS camera. ExF = excitation filter. EmF = emission filter. (B) A photograph of the Multiscope system constructed to image bacterial fluorescence (see Methods for full description of individual components (i) – (viii)). (C) Widefield fluorescence images of the bacteria sample mounted on agarose. Regions of interest which appeared in best focus were selected for processing. (D) Processed ROIs after K-means clustering. (E) Magnified image of (D). (F) Averaged fluorescence profiles for the top cluster in each K-means analysis. Profiles corresponding to first stimulation (black line) and second stimulation (red dashed line). Scale bars: (C) 500 µm (D) 100 µm (E) 50 µm.

A parallelised fluorescence microscope was built following the design of the Exeter Multiscope^12^, which in turn is based on the original RAP prototype^11^. Previous designs had been bright field only, detecting transmitted light with a machine vision camera. The key difference in this work was that we were detecting UV/VIS fluorescence and required UV excitation, and a scientific CMOS camera to achieve the required fluorescence sensitivity. Due to the in-line nature of the fluorescence detection, there was a need for strong filtering to block directly transmitted light, requiring the use of OD6 excitation (390 ± 20 nm) and emission (494 ± 10 nm) filters. The objective lenses had a diameter of 9 mm (limited by the sample pitch) and a focal length of 18 mm, providing an NA of 0.25 which was not sufficient to resolve individual bacteria, but bacterial microcolonies could be clearly discerned (Figure 2E).

The electrode array was loaded onto the sample tray and time lapse fluorescence images were acquired of *B. subtilis* samples at electrodes #1, #2, #5 and #6 (electrode positions #3 and #4 were not used as they were not centred on the 9 mm pitch of the objective lens array, Figure 1B). As previously, two sets of time lapse data were acquired for each electrode position, corresponding to two separate electrical stimulations to compare viable/non-viable responses. The Multiscope provided all-optical switching between the four fields of view by turning on the LED associated with each electrode position (Figure 2A). Only one field of view was imaged at any one time. An example widefield fluorescence image of electrode #2 is shown in Figure 2C. Within each 2048 x 2048 raw image, a smaller 256 x 256 pixel ROI was selected to reduce processing times for the K-means clustering analysis. ROIs were located where the bacterial colonies appeared in sharpest focus. To reduce the high level of background fluorescence, these ROI images were first processed using a rolling ball subtraction (50 pixel diameter) in FIJI^13^. The K-means clustering algorithm was then applied to the processed ROI data and each pixel assigned to a cluster (Figure 2D-E).

The time sequence of each pixel value in the ROI was then analysed using K-means clustering as for the inverted microscope data. The ‘top’ cluster (i.e. that most closely associated with the bacterial microcolony pixels) was then extracted. The peak difference in fluorescence intensity after stimulation was determined and the results are summarised in Table 2. The data set for electrode #6 was overwritten by the second stimulation and could not be analysed. To provide an indication of the signal to background ratio, Table 2*Table 2* also includes the mean value of each of the top cluster traces (‘signal’) and the bottom cluster traces (‘background’) where the bottom cluster corresponds to the pixels obscured by the opaque electrodes. The absolute change in fluorescence brightness appears to be approximately correlated with the signal to background ratio.

**Table 2:** Summary of fluorescence changes detected using the Multiscope at a bacteria density of OD=2.5. Bacteria at each electrode position were stimulated twice before moving to the next electrode.

| Electrode position | Stimulation number | Peak fluorescence change ($\Delta f/f$) | Top trace avg. (a.u.) | Bottom trace avg. (a.u.) | Signal to background |
| --- | --- | --- | --- | --- | --- |
| 1 | 1 | 4.0% | 61.1 | 7.9 | 7.7 |
| 1 | 2 | -6.7% | 60.3 | 7.9 | 7.6 |
| 2 | 1 | 13.9% | 74.0 | 8.7 | 8.5 |
| 2 | 2 | -10.7% | 74.5 | 8.7 | 8.8 |
| 5 | 1 | 9.4% | 126.4 | 9.2 | 13.7 |
| 5 | 2 | -10.6% | 120.3 | 8.9 | 13.5 |
| 6 | 2 | -4.7% | 99.5 | 9.3 | 10.6 |

### Multiscope data at lower cell densities

To further explore the sensitivity limits of the UV fluorescence Multiscope, we repeated the experiments at a lower density preparation of *B. subtills* (OD=1.5) for agarose pad inoculation. Reducing the bacterial density used for agarose pad inoculation offers additional clinical benefit by enabling earlier identification of effective antimicrobial compounds. Experiments were repeated as for the OD=2.5 samples. Multiscope widefield fluorescence images from each of the electrode positions are shown in Figure 3A, together with the selected ROI (red square). The processed data indicate that in some cases, such as electrode #1 (Figure 3B-C), the bacteria were very sparse.

**Figure 3.**
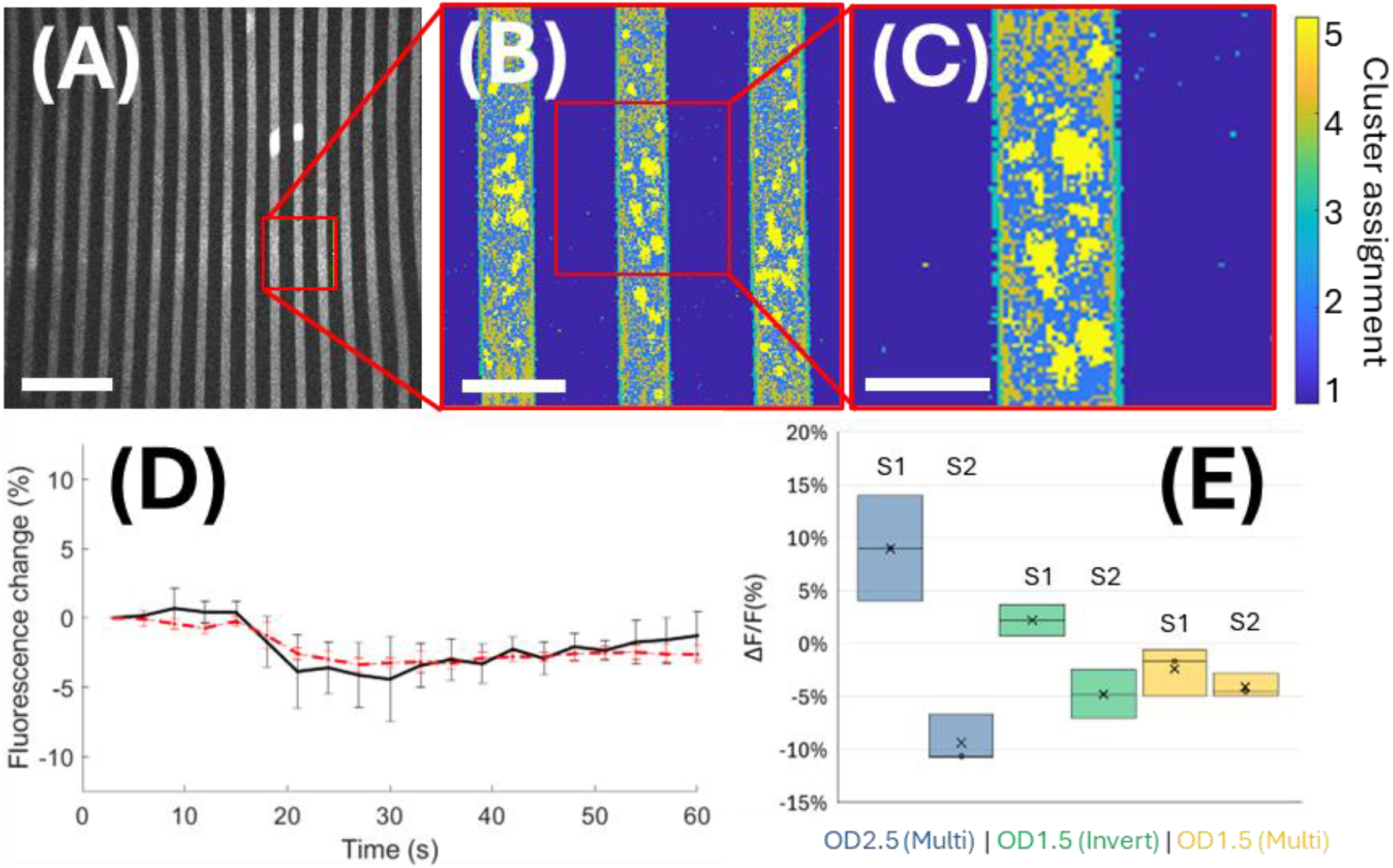
(A) Widefield fluorescence images of the bacteria samples mounted on agarose imaged through the opaque CytePulse electrodes. Regions of interest (red squares) were selected for processing. Example shown is a live cell response after the first stimulation. (B) Processed ROIs after K-means clustering. (C) Magnified view of (B). (D) Change in the fluorescence brightness for the top cluster averaged over three electrode positions after the first (black line) and second (red dashed line) stimulation. (E) Change in fluorescence brightness after first (S1) and second (S2) stimulation for the three conditions considered above (Multi=Multiscope, Invert=Inverted microscope).

The top cluster profiles are shown in Figure 3D. None of the traces indicate a significant change in fluorescence brightness. As shown in Table 3, the changes are all small with slightly negative changes in fluorescence. It is possible that, even though the signal to background levels are similar to that at high cell densities, there are simply not enough cells to register a significant change in fluorescence. Doubling the number of clusters from five to ten did not significantly affect the results. The more defocused images of the sample led to poorer rolling ball background subtraction leaving higher baseline levels of fluorescence for some electrode positions.

**Table 3:**
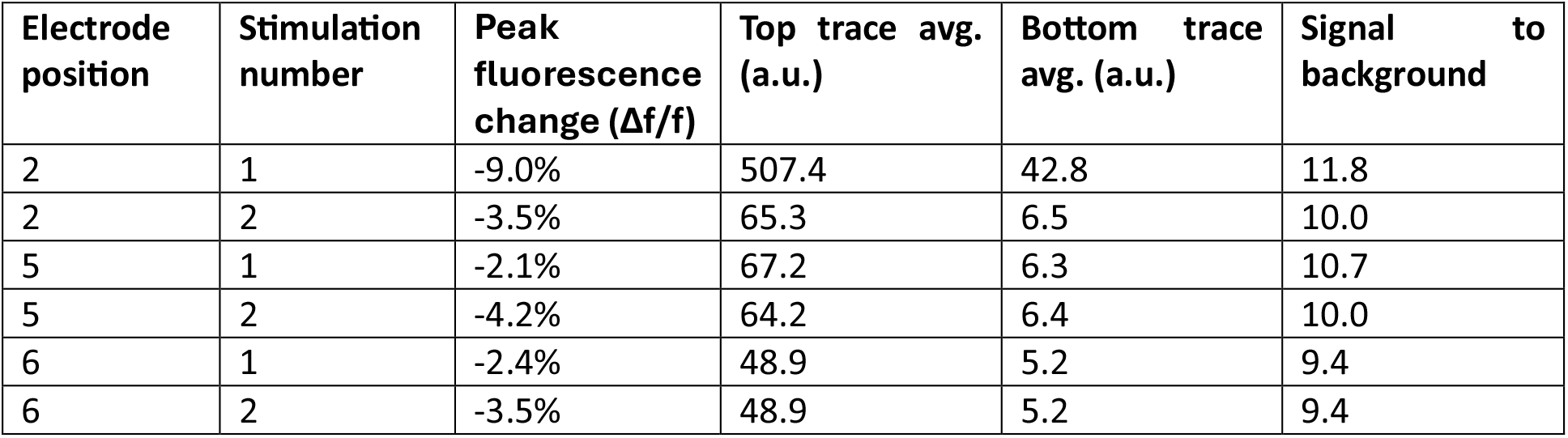
Measurements of peak fluorescence changes for each of the Multiscope experiments for a low bacteria density (OD=1.5).

## Discussion

Combining electrical stimulation with Multiscope readout enables the clear determination of viable from non-viable bacteria (Figure 2F). Differences in the amplitude of the fluorescence change (Figure 3E) are linked to both (i) contrast in the sample image and (ii) the strength of the fluorescence signal. Image contrast is maintained by keeping the sample in focus and minimising aberrations. Variations in sample focusing is thought to be largely due to the mounting of the electrode array in the Multiscope. Tension in the IDC cable produced a small uplift of electrode positions #2 and #5 relative to #1 and #6 (Figure 2B). Focusing the objective array on electrodes #2 and #5 led to reduced contrast in sample images at positions #1 and #6, as reflected in Table 2.

Aberrations from the parabolic reflector will also contribute to the reduction in image contrast. Comatic aberrations increase as *(B/f)*^2^ where *B* is the lateral distance of the sample from the optic axis of the mirror and *f* is the focal length of the mirror. This can be improved greatly by increasing the focal length of the mirror from the current value of 100 mm to around 300 mm and thereby reducing the comatic aberrations by a factor of nine. A longer parabolic mirror focal length would also increase the standoff distance of the sCMOS camera, reducing the number of sample positions that might be obscured by the camera itself.

The strength of the fluorescence signal has been shown to be dependent on both how well the sample was focused, and the abundance of bacterial colonies present in the sample, with ODs of 1.5 generating insufficient fluorescence signal. This could lead to the top cluster expanding to include more dim pixels from defocused bacterial microcolonies, thereby “averaging out” the strength of the fluorescence response. This is indicated by the number of top cluster pixels remaining approximately the same for the OD=1.5 samples (Figure 3B), despite the approximately two-fold reduction in bacterial abundance.

Future improvements in system throughput could be enabled by reducing the current 2.9 second integration time required to accumulate sufficient fluorescence signal from the bacterial monolayer. These long integration times undermine the benefits of all-optical switching between samples. Using UV sources that are brighter than the current 35 mW LEDs is one option. As only 1 mW of excitation light currently reaches the sample, there is significant potential to increase the optical coupling between source and sample by removing the plastic bulb covering the LED emitter and using higher NA collection lenses with higher transmission UV coatings. Transitioning from an in-line to an epi-fluorescent imaging architecture would also greatly increase contrast in the acquired images. The increase in signal to background ratio would also open up the possibility of obtaining reliable measures of viability at lower bacterial concentrations, even enabling imaging direct from biofluid, bringing the technology in line with competing AST approaches^2^.

If the camera integration time can be reduced from 2.9 seconds to around 500 ms (as used on the inverted microscope), then it would easily allow all four samples to be imaged in a single 45 second exposure by optically moving from one sample to the next between camera exposures. The raw image sequence could then be de-interlaced into four time-lapse sequences. If the integration time could be reduced further to a more typical 50 ms exposure, this would allow over 50 wells to be imaged simultaneously, including additional time budget for streaming images to disk. This is a significant target as it would allow for bacterial viability measurements in the presence of seven concentrations (plus controls) of the six main antibiotics administered clinically in acute infections (i.e. Piperacillin– tazobactam, Ceftriaxone, Gentamicin, Metronidazole, Vancomycin, Meropenem).

Finally, it can be seen in Figure 1C that the bacterial clusters were rarely distributed evenly across the agarose pad but would instead accumulate in a ‘coffee ring’ pattern left after the inoculation droplet had dried on the agarose pad. One potential modification to the protocol would be inoculating the agarose pad via spraying, as demonstrated in [^14^] to achieve a more uniform distribution across the pad.

## Conclusion

With the rapid emergence of AMR there is a growing and urgent need for rapid diagnostic methods capable of identifying antibiotic efficacy. Established ASTs are culture-based and inherently slow: even after a pure bacterial isolate has been obtained, standard AST methods typically require 24-48 hours of incubation to generate results. In contrast, bacterial optical electrophysiology can provide viability-dependent fluorescence readouts in under one minute, offering a fundamentally faster route for assessing antimicrobial effectiveness. To date, all previous implementations of the fluorescent measure of bacterial viability using electrical stimulation have used a standalone research grade microscope and require a high degree of user intervention to obtain bacterial viability data. In this work, we have demonstrated the viability of using an automated Multiscope approach to parallelise optical excitation and readout of fluorescence changes in bacterial monolayers. This provides an important stepping stone towards high throughput screening of bacterial populations using rapid optical electrophysiology methods. Whilst the initial test used a relatively modest number of testing sites (a 2 x 2 array) there is clear potential for scaling the number of tests to 50 or more wells using the same architecture, providing measures of antibiotic efficacy in one to two minutes. By matching rapid testing with high throughput, the goal of reducing bottlenecks in clinical AST can be achieved.

## Methods

### Multiscope construction

The prototype system in shown in Figure 2B. The light path starts with a 4 × 4 array of LEDs (RS Components #903-3771) which output up to 35 mW in the range 390-425 nm. As the light was emitted over a 30 degree cone angle, baffles were used to avoid light spilling over into adjacent well positions and creating cross-talk. The baffles were 3D printed as a 4 × 4 array of opaque tubes. These tubes were 6 mm in diameter and had a length which corresponded to the 9 mm focal length of the collimation lenses. The baffle was mounted onto the underside of the custom LED array and together occupied the first tray position (Figure 2B(i)). The second tray position was occupied by a 3 × 3 array of 9 mm diameter, 9 mm focal length collimation lenses (Edmund Optics, #65-549-INK). The short focal length was chosen to maximise the light transmitted onto the bacterial sample. By moving the lens array fractionally further away from the LED array, it was possible to focus the excitation light onto only the region of the bacterial sample that lies within the field of view of the objective lenses, further increasing the light intensity at the sample. A 50 mm diameter, OD6 excitation filter with a 390 ± 20 nm pass band (Edmund Optics #86-359) was placed after the collimation lenses. The filter was aligned to the well positions of the 4 × 4 LED array within a 3D printed mount that occupied the third well plate holder position (Figure 2B(iii)).

The CytePulse electrode array occupied the fourth tray position (Figure 2B(iv)). Directly underneath this layer was a 2 × 2 array of imaging objective lenses, centred on the samples (Figure 2B(v)). The four positions corresponded to electrode positions 1, 2, 5 and 6 (Figure 1(B)). These lenses (Edmund Optics #47-653, 9 mm diameter, 18 mm focal length, NA=0.25) collected the fluorescence emission from the bacteria and relayed this to the sCMOS camera (Hamamatsu ORCA Flash4.0 V2) via a parabolic mirror (100 mm focal length, 220 mm diameter, Edmund Optics #68-793). A 1” OD6 emission filter with pass band 494±10nm (Edmund Optics #84-096) was fixed immediately in front of the fluorescence camera to further suppress excitation light.

The 0.25 NA objective lenses provided both better light collection, higher spatial resolution and smaller depth of field to improve the sensitivity to fluorescence changes within the thin bacterial monolayer. The theoretical resolution of 1.2 µm combined with residual aberrations from the parabolic mirror meant it was not possible to resolve individual bacteria. The shallow depth of field required a finer degree of control over the objective lens position. This was provided by a single axis piezo (Figure 2B(vi), Mad City Labs MMP1) which achieved a minimum step size of 95 nm over a 25 mm range.

Fluorescence then reflected off the parabolic mirror (2B(vii)) and was collected by the sCMOS camera (Figure 2B(viii)). The magnification of the system is 100 mm / 18 mm = 5.5×. The maximum field of view of the system is therefore given by the size of the imaging chip (2048 × 6.5 um = 13.3 mm) divided by the magnification (13.3 mm / 5.5 = 2.4 mm square). To minimise stray light incident upon the camera, emanating from outside of the sample field of view, a 3D printed plate was used. This 2 mm thick plate contained an array of 3 mm diameter holes that acted as a field stop when centred on the CytePulse electrode locations and was aligned with the camera to avoid any clipping of the field of view.

### Data acquisition and hardware control

The Multi-dimensional acquisition toolbox in MicroManager^15^ was used to trigger the sCMOS camera and define the integration time. For the inverted microscope (IX73, Olympus), images were acquired on an sCMOS camera (Zyla 4.2, Andor) every second (500 ms exposure), on a 10× 0.4 N.A. dry objective. For the acquisition of Multiscope image data, a 2.9 second camera integration time on a 3 second interval was used for a total of 20 exposures. Electrical stimulation was manually triggered via the CytePulse graphical user interface 15 seconds after initiating image acquisition (i.e. after 15 frames on the inverted system and 5 frames on the Multiscope). Electrical stimulation of 4V amplitude and 100 Hz frequency for 2.5 seconds was used in all cases. All images in the sequence were saved directly to disk.

### Sample preparation

*B. Subtilis* 168 streak plates were prepared from cyrostock by streaking onto lysogeny broth (LB) agar plates and incubating overnight at 37 °C. Stationary-phase cultures were prepared by inoculating fresh LB with a single colony and incubating for 17 h at 37 °C with shaking. Log-phase cultures were prepared by diluting overnight cultures into fresh LB and incubating until mid-log phase (OD_600_ ≈ 1.5 or 2.5). Log-phase cells were used immediately for preparation of imaging samples.

A defined CytePulse medium (see Supplementary information) was prepared to support bacterial viability during imaging. MOPS and potassium phosphate buffers were adjusted to pH 7 and autoclaved, whereas trace-ion and carbon/nitrogen supplements (including amino acids, glucose, metal salts, and thiamine), and ThT were filter sterilised to avoid thermal degradation. ThT was included to enable fluorescence-based monitoring. Agarose pads (2% w/v) were prepared by combining molten agarose with the CyteCom medium.

Agarose pads were cast under aseptic conditions by pipetting molten agarose onto 22 × 22 mm coverslips and gently placing a second coverslip on top to form a uniform layer. After solidification, the upper coverslip was carefully removed, and each pad was cut into 16 equal sections with a sterile scalpel. Log-phase bacteria were spotted onto individual sections with a pipette and allowed to adsorb.

Each agarose section was then inverted onto a CytePulse electrode such that the bacterial layer contacted the electrode surface while avoiding the central ground electrode. The pads were then incubated at room temperature for 2 hours. Loaded electrodes were placed on a custom 3D-printed holders and transferred to either the inverted microscope or Multiscope for electrical stimulation and time-lapse imaging. This preparation yielded consistent single-layer bacterial samples suitable for combined CytePulse stimulation and fluorescence imaging.

## References

1. Interagency Coordination Group on Antimicrobial Resistance. No Time to Wait: Securing the Future from Drug-Resistant Infections. https://www.who.int/docs/default-source/documents/no-time-to-wait-securing-the-future-from-drug-resistant-infections-en.pdf (2019).

2. Reszetnik, G. et al. Next-generation rapid phenotypic antimicrobial susceptibility testing. Nat. Commun. 15, 9719 (2024).

3. Spencer, D. C. et al. A fast impedance-based antimicrobial susceptibility test. Nat. Commun. 11, 5328 (2020).

4. Stratford, J. P. et al. Electrically induced bacterial membrane-potential dynamics correspond to cellular proliferation capacity. Proc. Natl. Acad. Sci. 116, 9552–9557 (2019).

5. Cytecom Limited. https://www.cytecom.co.uk/.

6. Matsumoto, K. et al. Advanced CUBIC tissue clearing for whole-organ cell profiling. Nat. Protoc. 14, 3506–3537 (2019).

7. Gintoli, M. et al. Spinning disk-remote focusing microscopy. Biomed. Opt. Express 11, 2874 (2020).

8. Harfouche, M. et al. Imaging across multiple spatial scales with the multi-camera array microscope. Optica 10, 471 (2023).

9. Thomson, E. E. et al. Gigapixel imaging with a novel multi-camera array microscope. eLife 11, e74988 (2022).

10. Symvoulidis, P. et al. NeuBtracker—imaging neurobehavioral dynamics in freely behaving fish. Nat. Methods 14, 1079–1082 (2017).

11. Ashraf, M. et al. Random access parallel microscopy. eLife 10, e56426 (2021).

12. Mohanan, S. et al. Automated measurement of cardiomyocyte monolayer contraction using the Exeter Multiscope. Biomed. Opt. Express 16, 4716–4729 (2025).

13. Schneider, C. A., Rasband, W. S. & Eliceiri, K. W. NIH Image to ImageJ: 25 years of image analysis. Nat. Methods 9, 671–675 (2012).

14. Cano, Á. et al. Multiparametric quantification of bacterial cells using digital holographic microscopy. Sci. Rep. 15, 41051 (2025).

15. D. Edelstein, A. et al. Advanced methods of microscope control using μManager software. J. Biol. Methods 1, 1 (2014).

